# Sulcal organization in the medial frontal cortex reveals insights into primate brain evolution

**DOI:** 10.1101/527374

**Authors:** Céline Amiez, Jérome Sallet, William D. Hopkins, Adrien Meguerditchian, Fadila Hadj-Bouziane, Suliann Ben Hamed, Charles R.E. Wilson, Emmanuel Procyk, Michael Petrides

**Affiliations:** Univ Lyon, Université Lyon 1, Inserm, Stem Cell and Brain Research Institute U1208, 69500 Bron, France.; Wellcome Integrative Neuroimaging Centre, Department of Experimental Psychology, University of Oxford, Oxford OX1 3SR, United Kingdom.; Department of Comparative Medicine, University of Texas MD Anderson Cancer Center, Bastrop, Texas, 78602, USA.; Laboratoire de Psychologie Cognitive, UMR7290, Université Aix-Marseille, CNRS, 13331 Marseille, France.; Station de Primatologie CNRS, UPS846, 13790 Rousset, France.; Brain & Language Research Institute, Université Aix-Marseille, CNRS, 13604 Aix-en-Provence, France.; Integrative Multisensory Perception Action & Cognition Team (ImpAct), INSERM U1028, CNRS UMR5292, Lyon Neuroscience Research Center (CRNL), Lyon, France; University of Lyon 1, Lyon, France.; Institut des Sciences Cognitives Marc Jeannerod, UMR5229, CNRS-Université Claude Bernard Lyon I, 67 Boulevard Pinel, 69675 Bron, France.; Montreal Neurological Institute, Department of Neurology and Neurosurgery and Department of Psychology, McGill University, Montreal, Quebec, Canada.

## Abstract

Although the relative expansion of the frontal cortex in primate evolution is generally accepted, the nature of the human uniqueness, if any, and between-species anatomo-functional comparisons of the frontal areas remain controversial. To provide a novel interpretation of the evolution of primate brains, sulcal morphological variability of the medial frontal cortex was assessed in old-world monkeys (macaque, baboon) and Hominoidea (chimpanzee, human). We discovered that both Hominoidea do possess a paracingulate sulcus, which was previously thought to be uniquely human and linked to higher cognitive functions like mentalizing. Also, we revealed systematic sulcal morphological organisations of the medial frontal cortex that can be traced from multiple old-world monkey to Hominoidea species, demonstrating an evolutionary conserved organizational principle. Our data provide a new framework to compare sulcal morphology, cytoarchitectonic areal distribution, connectivity, and function across the primate order, leading to clear predictions on how other primate brains might be anatomo-functionally organized.

## INTRODUCTION

Is the human brain unique? Although the relative expansion of the frontal cortex in primate evolution is generally accepted, the nature of the changes that occur remains controversial. Neuroanatomy offers a window into the evolution of brain circuits and its correlates, brain functions. Looking at a macroscopic level, a large literature has emphasized the link between the extent of gyrification, the rapid expansion of the cerebral cortex, and the complexity of the computational processing performed in a given brain (e.g. Zilles et al., 1989, 2013). Several hypotheses have been proposed to explain the origin of sulci and gyri: genetic control (Rakic, 2004), cortical growth (Toro and Burnod, 2005), tension of white matter cortico-cortical axons (Van Essen, 1997; Hilgetag and Barbas, 2005, 2006), cortico-thalamic axons (Rakic, 1988), or simply mechanical instability in a soft tissue growing non-uniformly (Tallinen et al., 2014; Amiez et al., 2018; Van Essen et al., 2018). Although important, these discussions of cortical gyrification have not considered another major dimension of sulcal pattern organization, i.e. its variability. Interindividual variability of traits are at the basis of many evolutionary genetic studies (Harris, 1966; Przeworski et al., 2000). Therefore, the primary goal of our study was to investigate variability in sulcal phenotypes from four key primate populations to provide a new understanding into evolution of primate frontal cortex.

Sulcal organization is not random despite the presence of strong inter-hemispheric and inter-subject variability (Petrides, 2012). Rather, it follows a precise topographical organization. The most straightforward example is the location of the central sulcus. Although its shape, length, and depth may vary across hemispheres and individuals, it is always present and systematically located at the same strategic antero-posterior location. Importantly, this is not a human specific feature as it is observed in all primates (Sherwood et al., 2004; Hopkins et al., 2014). In primates, although the origin of gyrification is not well understood, three types of sulci can be identified based on their appearance during gestation: primary sulci (e.g. central sulcus) are the earliest formed during gestation (i.e. before the 30^th^ week of gestation in human), followed by secondary sulci (e.g. paracingulate sulcus, formed between the 32^nd^ and the 36^th^ week of gestation in human), and finally tertiary sulci (formed between the 36^th^ week of gestation and the end of the first year of life in human) (Armstrong et al., 1995; Tamraz and Comair, 2000). Importantly, although primary sulci are present in all hemispheres in all individuals, the probability of observing secondary/tertiary sulci is variable. For instance, the cingulate sulcus (CGS) in the Medial Frontal Cortex (MFC) is a primary sulcus and as such is observed in 100% of hemispheres; by contrast the paracingulate sulcus (PCGS), a sulcus running parallel to the CGS, is a secondary sulcus and is present only in about 70% of human subjects at least in one hemisphere (Vogt et al., 1995, 2005; Paus et al., 1996; Amiez et al., 2018).

Importantly, several lines of evidence indicate that primary sulci are limiting sulci between cytoarchitectonic areas. For instance, the central sulcus is the limiting sulcus between the primary motor cortex (area 4) and the primary somatosensory cortex (area 3). In addition, one study (Vogt et al., 2005) has shown that the cytoarchitectonic organization of the cingulate cortical region is modulated, depending on the presence/absence of a secondary sulcus in the cingulate cortex, namely the PCGS. In addition, several studies performing single subject analysis have demonstrated that the organization of sulci and its variability is also pertinent to prediction of the localization of functional areas (Tomaiuolo and Giordano, 2016). For instance, functional magnetic resonance imaging (fMRI) studies have shown that the primary hand motor area is always located in the central sulcus in human and non-human primates (Yousry et al., 1997; Amiez and Petrides, 2014, 2018; Hopkins et al., 2014), the frontal eye field is always located in the ventral branch of the superior precentral sulcus in the human brain (Amiez et al., 2006; Amiez and Petrides, 2009, 2018; Derrfuss et al., 2012) and in the genu of the arcuate sulcus in macaque (Bruce and Goldberg, 1985; Koyama et al., 2004), the face motor representation of the rostral cingulate motor area and the juice feedback-related activity during exploratory situations recruits a region located in the CGS when no PCGS is present, but in the PCGS when the CGS is present (Amiez et al., 2013; Amiez and Petrides, 2014). Comparable observations were made in other regions of the brain, such as the inferior frontal junction (Derrfuss et al., 2009, 2012), the orbitofrontal cortex (Li et al., 2015), the parietal somatomotor cortex (Zlatkina et al., 2016), and the angular gyrus region (Segal and Petrides, 2013). Collectively, these studies strongly suggest that the sulcal organization in primates is not random, but rather has anatomo-functional relevance that are likely associated with the evolution to increasingly sophisticated sensory, motor, and cognitive functions. To what extent sulcal morphology across primate brains can provide insights into the anatomo-functional organization of the brains in higher primates and brain evolution in the primate order is, however, poorly understood.

The aim of the present study was to determine how the MFC sulcal organization has evolved through the primate order, a region often neglected in comparative studies (Van Essen and Dierker, 2007), despite its key role in cognitive functions often thought to be uniquely human, such as mentalizing or counterfactual thinking (Amodio and Frith, 2006). Opposing views suggest that the MFC is functionally comparable in humans and macaque (Sallet et al., 2013; Amiez et al., 2016). As mentioned above, several studies have shown that strong relationships exist between sulcal morphology and functional organization in the human brain. Based on neuroimaging anatomical scans, we investigated the sulci morphological variability from old-world monkeys (80 macaques and 88 baboons) and Hominoidea (225 chimpanzees and 197 humans) to identify potential specific properties of the human MFC. We discovered that the analysis of the sulcal organization in the MFC in primates provides critical new evidence of how the prefrontal cortex evolved within the order Primates. We discovered that both Hominoidea do possess a paracingulate sulcus, which was previously thought to be uniquely human and linked to higher cognitive functions like mentalizing. We also confirmed the expansion of the most rostral part of the medial prefrontal cortex in Humans. But overall, we revealed systematic sulcal morphological organisations of the medial frontal cortex that can be traced from multiple old-world monkey to Hominoidea species, demonstrating an evolutionary conserved organizational principle.

## RESULTS

### The PCGS, an ape innovation

A major finding of the present analysis was the demonstration of the presence of the PCGS in at least one hemisphere within human and chimpanzee brains but not in baboon and macaque brains (χ2 = 242.18, df = 3, p-value = 2.2e-16, logistic regression, GLM fitted using an adjusted-score approach to bias reduction) (Fig 1A, B, C, D, E).

**Figure 1.**
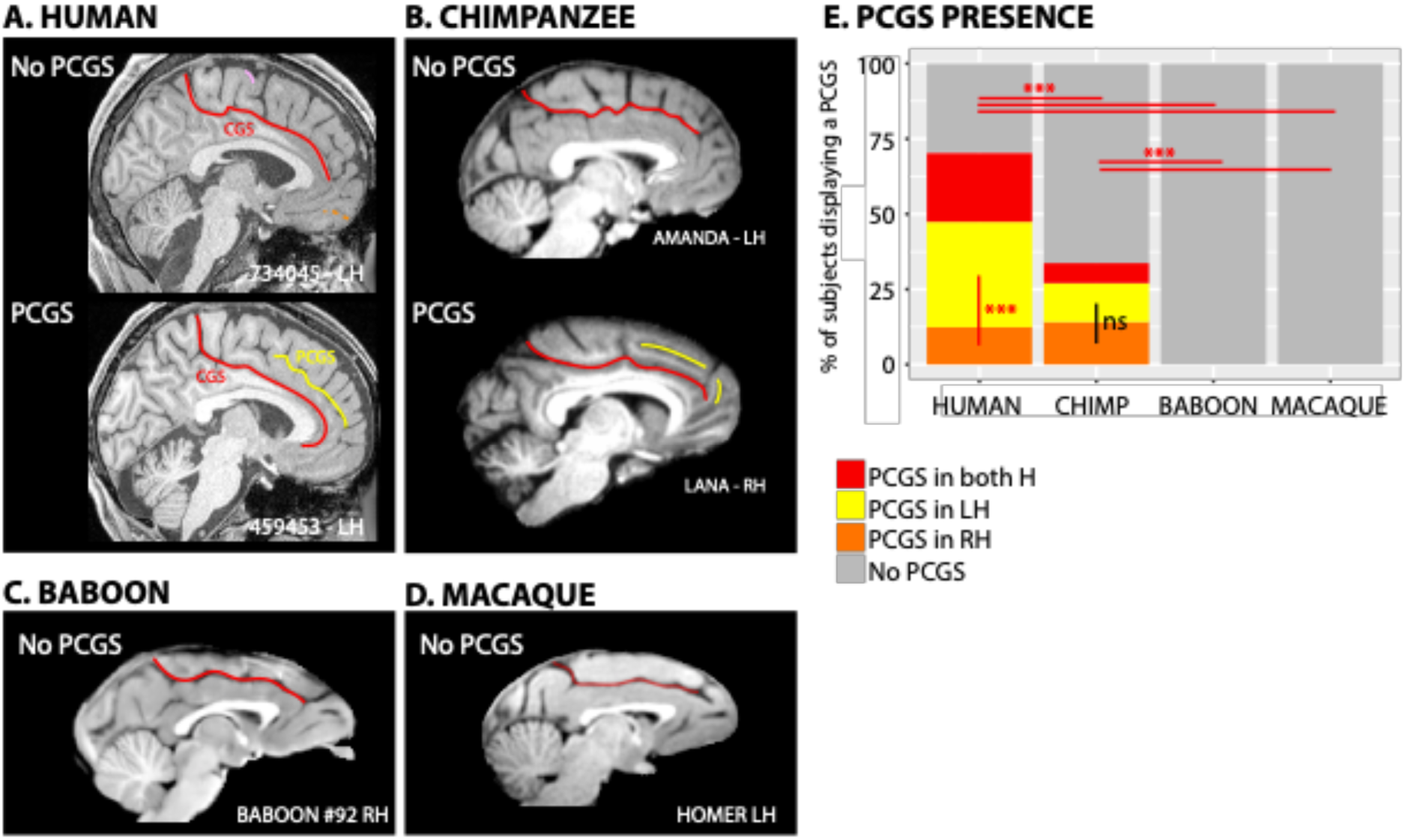
Presence of the paracingulate sulcus (PCGS) across primates. The location of the cingulate sulcus (CGS) is shown in red in typical hemispheres displaying no PCGS versus those displaying a PCGS in human (A), chimpanzee (B), and in typical hemispheres of baboon (C) and macaque (D) brains. The location of the PCGS is shown in yellow in typical hemispheres displaying a PCGS in human (A) and chimpanzee (B). The present analysis demonstrated the presence of PCGS in both human and chimpanzee brains, but not in baboon and macaque brains (E). The probability of occurrence of a PCGS decreased from human to chimpanzee. The PCGS is lateralized in the left hemisphere in human, but not in chimpanzee, brains. Statistics: * p<0.05, ** p<0.01, *** p<0.001, ns non-significant.

A post-hoc Tukey test revealed that the probability of occurrence of a PCGS differed significantly between human and chimpanzee brains with a PCGS in least in one hemisphere in 70.1% of human versus 33.8% of chimpanzee brains (estimate=1.47562, std. error=0.20917, value=7.054, p-value<0.001) (Fig 1E). Interestingly, humans and chimpanzees differed with respect to asymmetries in the PCGS. In human brains, the PCGS was present in the left hemisphere only in 35% of subjects, in the right hemisphere only in 12.7% of subjects, and in both hemispheres in 22.8% of subjects, a distribution that differed significantly (χ2 = 19.912, df = 1, p-value = 8.109e-06, logistic regression) (Fig 1E). In contrast, in chimpanzee, the PCGS was present in the left hemisphere only in 12.9% of subjects, in the right hemisphere only in 14.2% of subjects, and in both hemispheres in 6.7% of subjects, a distribution that did not significantly differ (χ2 = 0.12399, df = 1, p-value = 0.7247).

The presence or absence of an intralimbic sulcus (ILS, i.e. a shallow sulcus running parallel and ventral to the cingulate sulcus), was also characterized in our primate sample. The ILS was observed in at least one hemisphere in 27%, 18.7%, 12.5%, and 11.3% of human, chimpanzee, baboon, and macaque brains (Fig 2A), which were significantly different (χ2 = 18.993, df = 3, p-value = 0.0002743, logistic regression). Post-hoc Tukey analysis revealed that the presence of ILS in humans was significantly higher than all other primate species. Further, no significant differences in the presence of the ILS were found between the three non-human primate species (chimpanzee versus baboon: p=0.5502, chimpanzee versus macaque: p=0.4176, baboon versus macaque: p=0.9942).

**Figure 2.**
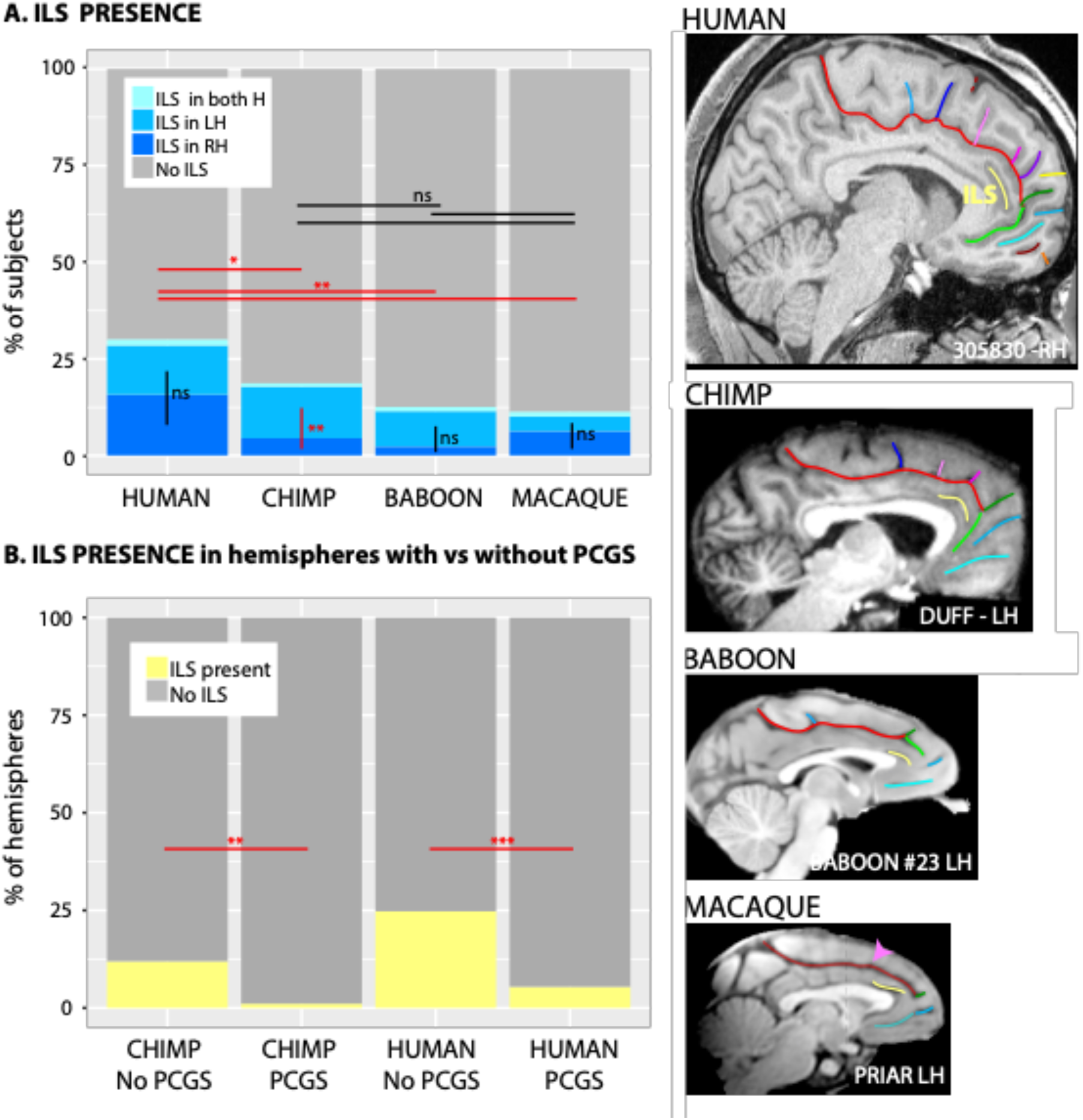
Presence of the intralimbic sulcus (ILS) across primates. The location of the ILS, i.e. a shallow sulcus running parallel and ventral to the cingulate sulcus, is shown in yellow in typical hemispheres of human, chimpanzee, baboon, and macaque brains (right panels). **A**. The probability of occurrence of an ILS is higher in human brains compared with non-human primates. A post-hoc Tukey test indicated no significant difference in the presence of the ILS in the three non-human primate species. Only in chimpanzee brains, the probability to observe an ILS is higher in the left than in the right hemisphere. By contrast, no difference between the f probability of occurrence of ILS in the left versus right hemisphere was found in human, baboon, and macaque brains. **B**. in Hominoidea, when a PCGS is present, the ILS is almost absent. Statistics: * p<0.05, ** p<0.01, *** p<0.001, ns non-significant.

We also tested for asymmetries in the presence of ILS within each species. No interhemispheric differences were found in humans (χ2 = 0.68999, df = 1, p-value = 0.4062 (ns), baboons (χ2 = 3.3591, df = 1, p-value = 0.06684 (ns), or rhesus monkeys (χ2 = 0.42938, df = 1, p-value = 0.5123 (ns), logistic regression). However, the ILS was found to be present significantly more often in the left than right hemisphere in chimpanzees (χ2 = 10.419, df = 1, p-value = 0.001247, logistic regression). Lastly, in Hominoidea, when a PCGS was present, the ILS was almost always absent (Fig 2B) (Human: presence of ILS in hemispheres with versus hemispheres without a PCGS: χ2 = 30.221, df = 1, p-value = 3.855e-08; Chimpanzee: χ2 = 14.011, df = 1, p-value = 0.0001817, logistic regression).

### The dorsal medial frontal cortex, a well-preserved structural organization across primates

There are several lines of evidences suggesting that the dorsal medial frontal cortex is comparable in terms of cytoarchitectonic areal distribution, connectivity, and functions across macaque and human brains (Cole et al., 2009, 2010; Procyk et al., 2016; Vogt, 2016). The present analysis adds up on these results revealing that the sulcal organization of this region is well-preserved too.

In the human brain, posterior to the genu of the corpus callosum, we identified four sulci vertical to the CGS or the PCGS including the paracentral sulcus (PACS), the pre-paracentral sulcus (PRPACS), the posterior (VPCGS-P) and the anterior (VPCGS-A) vertical paracingulate sulci (Fig 3). For each of these sulci, we characterized them as either fully present, as a spur or dimple when only superficially evident, or absent. For this analysis, we considered the presence of spurs or dimples, which were observed almost entirely in the baboon and rhesus monkey samples, as precursors to their full sulci expression in chimpanzees and humans (see below and FIG S1). Our argument for considering spurs and dimples as precursors of each of the 4 sulci in rhesus monkeys and baboons comes from an analysis of their normalized spatial location relatively to human and chimpanzee brains (see below). Specifically, we observed that the PACS, PRPACS, VPCGS-P and VPCGS-A sulci, spurs or dimples could be reliably located with respect to four anatomical landmarks: the rostral limit of the pons, anterior commissure, caudal limit of the genu of the corpus callosum, and the rostral limit of corpus callosum (see FIG 3A). To assess the consistency in the occurrence of PACS, PRPACS, VPCGS-P and VPCGS-A with respect to these landmarks, we calculated the difference between the Y value of the intersection between the CGS and the PACS and the Y value of each anatomical landmark. Note that the MRI data of the four species were first normalized in their respective normalized space (see Method). This difference was then normalized to take into account the different antero-posterior extent of the brains of the four species (i.e. antero-posterior extent of the human, chimpanzee, baboon, and macaque brains are respectively 175, 110, 85 and 60 mm). These results are shown in FIG3B to 3E and, as can be seen, the variability around the standardized location of each sulcus relative to their anatomical landmark were consistent within and between species.

**Figure 3.**
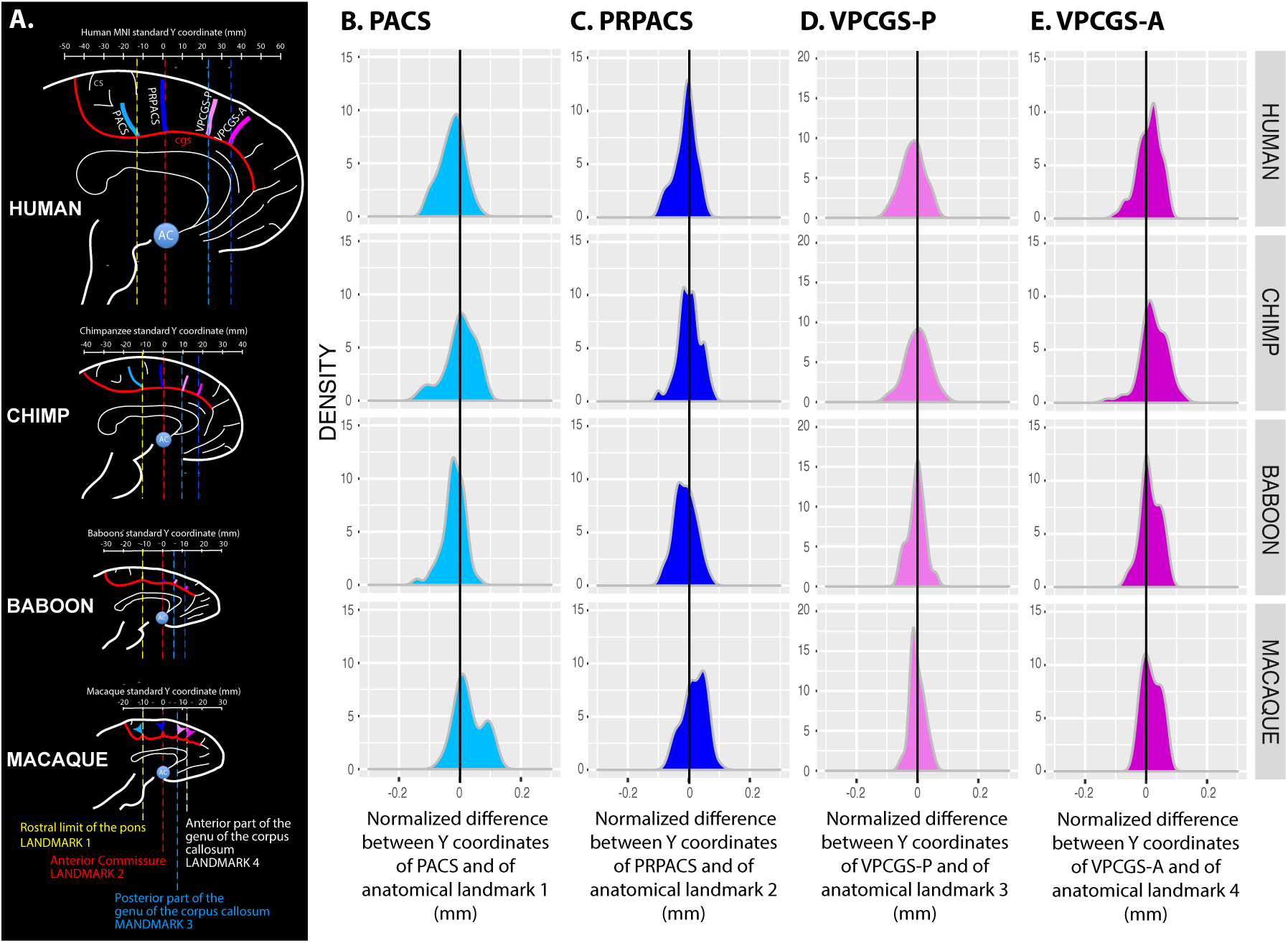
Location of vertical sulci in the dorsomedial frontal cortex across primates. **A**. Schematic representation of the location of the various sulci in the respective standard space of each primate species. The scale represents the antero-posterior level (in mm) in each brain. Anatomical landmarks fixed across primates are displayed: the rostral limit of the pons (landmark 1, in yellow), the anterior commissure (landmark 2, in red), the posterior limit of the genu of the corpus callosum (landmark 3, in light blue), and the anterior limit of the genu of the corpus callosum (landmark 4, in dark blue). **B**. Density of the difference between the Y coordinates of PACS and of the anatomical landmark 1 in human, chimpanzee, baboon, and macaque (from top to bottom panels). This difference was normalized in relation to the brain size (total antero-posterior extent). The 0 value correspond to the Y level of the anatomical landmark 1 in standard brains. **C**. Density of the normalized difference between the Y coordinates of PREACS and of the anatomical landmark 2. The 0 value correspond to the Y level of the anatomical landmark 2 in standard brains. **D**. Density of the normalized difference between the Y coordinates of P-VPCGS and of the anatomical landmark 3. The 0 value correspond to the Y level of the anatomical landmark 3 in standard brains. **E**. Density of the normalized difference between the Y coordinates of A-VPCGS and of the anatomical landmark 4. The 0 value correspond to the Y level of the anatomical landmark 4 in standard brains.

When evaluating the probability of occurrence of each of the 4 vertically oriented sulci along the CGS or PCGS, significant species differences were found for PACS (χ2 = 43.068, df = 1, p = 2.381e-09), PRPACS (χ2 = 49.192, df = 3, p = 1.187e-10), VPCGS-P (χ2 = 108.8, df = 3, p = 2.2e-16) and VPCGS-A (χ2 = 83.311, df = 3, p = 2.2e-16) (see FIG 3A to 3D). For PACS, VPCGS-P and VPCGS-A, post-hoc analysis indicated that these sulci were present in a higher proportion of human brains compared to chimpanzees, rhesus monkeys and baboons. Additionally, the percentage of chimpanzee brains displaying a PACS, VPCGS-A and VPCGS-P was significantly higher compared to rhesus monkeys and baboons; however, there was no significant difference in the probability of occurrence between baboons and rhesus monkeys. For PRPACS, post-hoc analysis indicated that its probability of occurrence was significantly higher in humans and chimpanzee brains compared to rhesus monkeys and baboons. There were no significant differences in its occurrence between human and chimpanzee brains nor between baboon and macaque brains.

In sum, the four vertical sulci, or their precursors, can be found across the four primate species examined in the current study. In humans, the four sulci are equally and largely present in both hemispheres (χ2 = 4.6815, df = 3, p = 0.1967). By contrast, their co-occurrence is not equally present in chimpanzees (χ2 = 68.917, df = 1, p = 7.279e-15), baboons (χ2 = 38.617, df = 1, p = 2.092e-08), and rhesus monkeys (χ2 = 67.617, df = 1, p = 1.382e-14]. As can be seen in Fig 4A to D, PACS, PRPACS, and VPCGS-A are the most conserved sulci across primate species, while VPCGS-P is the least conserved sulcus (Fig 4). Chimpanzee brains display a sulcal organization of the dorsal medial frontal cortex comparable to that of the human brain, macaque and baboon brains display evidence of the emergence of this organization. Importantly, vertical sulci emerging from the CGS or the PCGS located in the dorsal medial frontal cortex can be identified within fixed anatomical landmarks that are present across primates.

**Figure 4.**
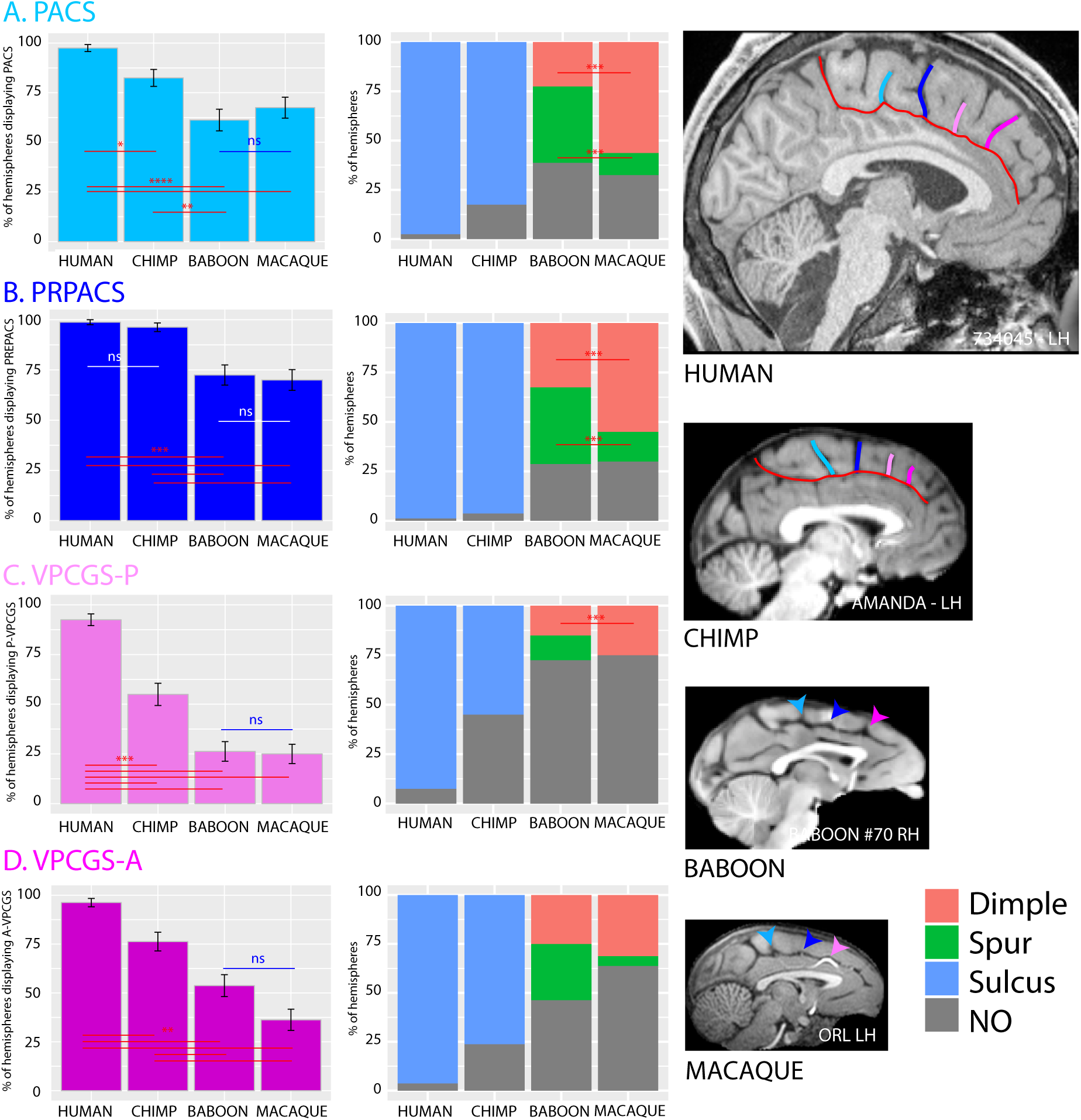
Presence of vertical sulci in the dorsomedial frontal cortex through primates. Probability of occurrence of the paracentral sulcus (PACS, A), the pre-paracentral sulcus (PRPACS, B), the posterior (VPCGS-P, C) and the anterior (VPCGS-A, D) vertical sulcus in the four species. **A. PACS. Left panel:** The probability to observe a PACS decreases from human, chimpanzee, baboon, to macaque. Posthoc analysis revealed that the PACS presence was higher in Hominoidea than in Cercopithecinae but that is was similar between human and chimpanzee on one hand, and between baboon and macaque on the other hand. **Right panel:** PACS characteristics varies across primate: it is a sulcus (blue areas) in 100% of hemispheres in Hominoidea (human and chimpanzee), it is a spur (green areas) or a dimple (pink areas) in Cercopithecinae (baboon and macaque). It is more frequently a spur than a dimple in baboon, as opposed to macaque. **B. PRPACS. Left panel:** the probability to observe PACS is higher in Hominoidea than in Cercopithecinae but is similar between human and chimpanzee on one hand, and between baboon and macaque on the other hand. **Right panel:** PRPACS characteristics varies across primate: it is a sulcus in 100% of hemispheres in Hominoidea (human and chimpanzee), it is a spur or a dimple in Cercopithecinae (baboon and macaque). It is more frequently a spur than a dimple in baboon, as opposed to macaque. **C. VPCGS-P. Left panel:** the probability to observe VPCGS-P decreases from human, chimpanzee, baboon, to macaque. Posthoc analysis revealed that the VPCGS-P presence was higher in human than in non-human primates. Its probability of occurrence was also higher in chimpanzee than in Cercopithecinae. It was, however, similar between macaque and baboon. **Right panel:** VPCGS-P characteristics vary across primates: it is a sulcus in 100% of hemispheres in Hominoidea (human and chimpanzee), it is a spur or a dimple in baboon and only a dimple in macaque. **D. VPCGS-A. Left panel:** The probability of occurrence of VPCGS-A decreases from human, chimpanzee, baboon, to macaque. **Right panel:** VPCGS-P characteristics vary across primates: it is a sulcus in 100% of hemispheres in Hominoidea (human and chimpanzee), it is more frequently a spur than a dimple in baboon, as compared with macaque. Statistics: * p<0.05, ** p<0.01, *** p<0.001, ns non-significant.

### The anterior cingulate (ACC), ventral medial prefrontal (vmPFC) and the frontopolar cortex

The ACC, vmPFC, and medial frontopolar cortex are the focus of many studies on, respectively, emotional processing (Caruana et al., 2018), value-based decision-making (Rushworth et al., 2011; Papageorgiou et al., 2017), and high-order socio-cognitive processing (e.g. mentalizing (Amodio and Frith, 2006; Vaccaro and Fleming, 2018; Wittmann et al., 2018)). These abilities reach their epitope in humans, strongly suggesting that these cortical areas have evolved. The present analysis reveals how these regions have evolved, providing a structural substrate for improved abilities in the primate order.

Anterior to the genu of the corpus callosum, we identified in human brains up to 9 distinct sulci (Lopez-Persem et al. in revision). One of the distinct features of the vmPFC is the branching of sulci at the end of the CGS. In human brains, the rostral end of the cingulate sulcus is characterized by two sulci forming a downward bifurcation: the supra-rostral sulcus (SU-ROS) and the supra-orbital sulcus (SOS) (Fig 5A). When the PCGS is absent, the fork is located at the rostral end of the CGS but, when the PCGS is present, it is located at the rostral end of the PCGS in the large majority of hemispheres (Fig S2).

**Figure 5.**
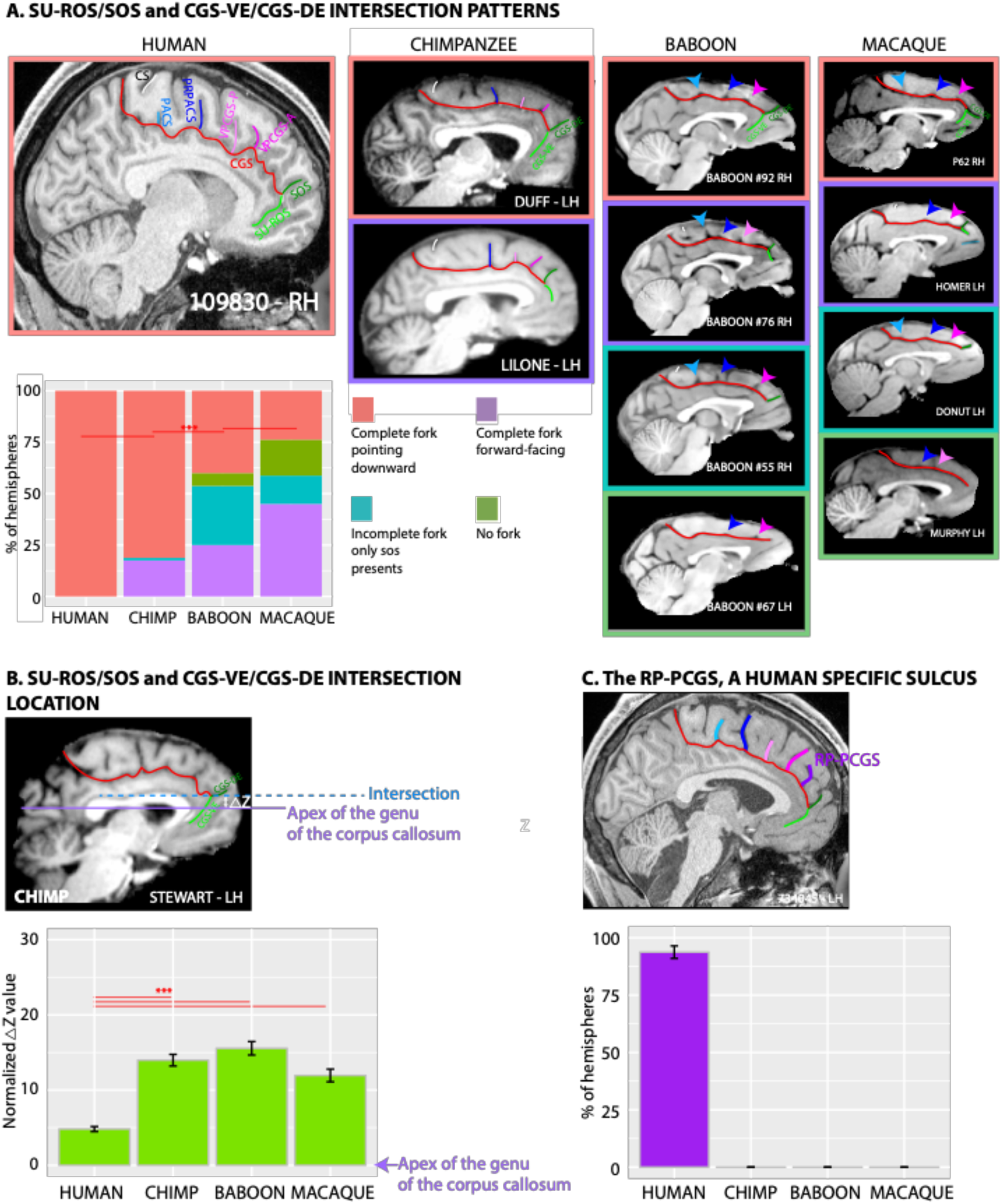
Morphological characteristics of the junction between the dorsal and ventral medial frontal cortex across primates. A. Rostral end of the CGS/PCGS. **In Human.** The rostral end of CGS is characterized by two sulci forming a fork pointing downward (pink area): the supra-rostral sulcus (SU-ROS) and the sus-orbitalis sulcus (SOS). **In chimpanzee**. The fork formed by the precursors of SU-ROS and SOS, i.e. the ventral (CGS-VE) and dorsal (CGS-DE) extensions of the cingulate sulcus, respectively, are also pointing downward in the majority of hemispheres. In 17.5% of hemispheres, the fork is forward facing (purple area), and is incomplete (green area) (i.e. SOS is present but SU-ROS sulcus is absent) in 1.25% of hemispheres. **In baboon**. The fork formed by CGS-VE and CGS-DE is also pointing downward in the majority of hemispheres but to a lesser extent than in Hominoidea. In 25% of hemispheres, the fork is forward facing, it is incomplete in 28.75% of hemispheres, and is absent in 6.25% of hemispheres. **In macaque**. The fork formed by CGS-VE and CGS-DE is displaying the same possible patterns than the baboon but in different proportions: the most frequent pattern observed is a fork facing forward, followed by a fork looking downward. The fork is incomplete in 13.75% of hemispheres and is absent in 17.5% of hemispheres. Statistics: * p<0.05, ** p<0.01, *** p<0.001, ns non-significant. **B. Location of the SU-ROS/SOS and CGS-DE/CGS-VE intersection**. The right diagram shows the location of the intersection between SU-ROS/SOS and CGS-VE/CGS-DE in a typical hemisphere of a chimpanzee. The ΔZ correspond to the difference between the dorsoventral Z coordinates where the intersection is observed and the dorsoventral Z coordinates where the apex of the genu of the corpus callosum is located. ΔZ was then normalized for brain size to compare across primate brains. Mean normalized ΔZ across individuals is displayed in the left diagram. The figure shows that the intersection of this intersection with the CGS or the PCGS is located close to the level of the apex of the genu of the corpus callosum, but dorsal to it in non-human primates. **C. Probability of occurrence of RP-PCGS**. The RP-PCGS is human-specific as non-human primates do not display it. Statistics: * p<0.05, ** p<0.01, *** p<0.001, ns non-significant.

In the non-human primates, a majority of the brains displayed different sulcal patterns at the rostral end of the CGS. The complexity and orientation of these folds were highly variable between species. Because of this discrepancy with the observed pattern in human brains, we chose to label the sulci at the end of the CGS, the ventral (CGS-VE) and dorsal (CGS-DE) extension of the cingulate sulcus. We viewed the CGS-VE and the CGS-DE in the nonhuman primate brains as precursors of the human SU-ROS and SOS respectively.

In chimpanzees, the fork formed by the CGS-VE and CGS-DE extension of the cingulate sulcus was also oriented downward in the majority of hemispheres (81.25%). However, in 17.5% of hemispheres, the fork was forward facing, and was incomplete (i.e. SOS is present but SU-ROS sulcus is absent) in 1.25% of hemispheres (Fig. 5A). Interestingly, as in humans, when the PCGS is absent, the fork is located at the rostral end of the CGS. By contrast, when the PCGS is present in the chimpanzee brain, the fork was is still located at the rostral end of the CGS -and not at the PCGS as in human-in the large majority of hemispheres (Fig S2).

In baboons, the fork formed by CGS-VE and CGS-DE is also oriented downward in the majority of hemispheres but to a lesser extent (40%) compared to humans and chimpanzees. In 25% of hemispheres, the fork is forward facing, is incomplete in 28.75% of hemispheres, and absent in 6.25% of hemispheres (Fig. 5A). In rhesus monkeys, the fork formed by CGS-VE and CGS-DE displays the same patterns than in baboons but in different proportions: the most frequent pattern observed is a forward-facing fork (in 45% of hemispheres), followed by a fork oriented downward (in 23.75% of hemispheres). The fork is incomplete in 13.75% of hemispheres and is absent in 17.5% of hemispheres (Fig. 5A).

To assess the relative location of the intersection of this fork with the CGS or PCGS across primates, we calculated the percentage of displacement of this intersection from the apex of the genu of the corpus callosum. This displacement was normalized across primates on the basis of the dorso-ventral extent of the brains normalized in their respective standard space (see Methods) at the level of the genu of the corpus callous across primates (i.e. 100mm, 50mm, 30mm, and 24mm, respectively in human, chimpanzee, baboon, and macaque brains). Results showed that the intersection of this fork with the CGS or the PCGS is located at the level of the apex of the genu of the corpus callosum in humans (Fig 5B), but is located dorsal to it in the non-human primate brains (F_(3,298)_=43.506, p<2.2e-16, GLM), strongly suggesting that this has migrated downward in the human brain.

Ventral to SU-ROS or CGS-VE lies the superior rostral sulcus (ROS-S). The ROS-S is a long and deep sulcus that is present in nearly 100% of hemispheres across all four species and therefore does no differ significantly in its occurrence (χ2 = 4.5334, df = 3, p = 0.2093) (Fig 6A). The ROS-S is often connected to the accessory supra-orbital sulcus (ASOS) and this sulcus is equally present in human (88.75%) and chimpanzee (81.25%) hemispheres. The ASOS is observed significantly less often in the baboon (26.25%) and rhesus monkey (21.25%) hemispheres compared to the chimpanzees and humans (χ2 = 132.81, df = 3, p-value = 2.2e-16, logistic regression (Fig 6B).

**Figure 6.**
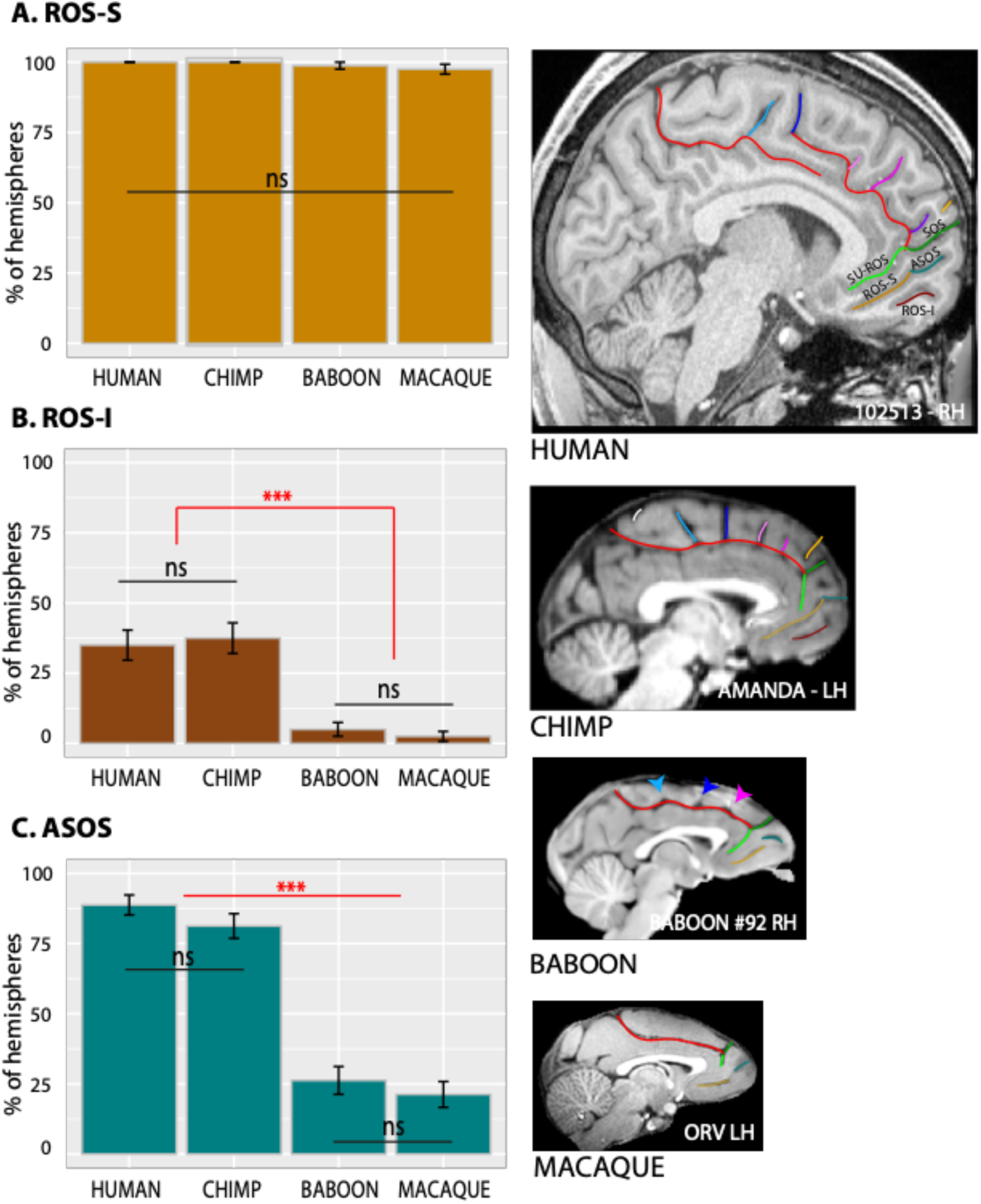
Sulci present in the ventral medial frontal cortex through primates. The location of the ROS-S, ROS-I, and ASOS sulci are displayed in human, chimpanzee, baboon, and macaque brains in the right panels. **A**. The probability of occurrence of ROS-R is similar in human and non-human primates. **B**. The probability of occurrence of ROS-I is identical in human and chimpanzee and is almost absent in Cercopithecinae. **C**. The probability of occurrence of ASOS is equal in human and chimpanzee. It is decreased in Cercopithecinae but equally present in baboon and macaque. Statistics: * p<0.05, ** p<0.01, *** p<0.001, ns non-significant.

The most ventral sulcus within the vmPFC is the inferior rostral sulcus (ROS-I). This sulcus is equally present in human (35%) and chimpanzee (37.5%) hemispheres but largely absent in the baboons (3%) and rhesus monkeys (1.5%) (χ2 = 60.348, df = 3, p = 4.952e-13,) (Fig 6C).

Finally, we observed at the most rostral part of the frontal cortex, several sulci that are almost exclusively present in Hominoidea. Notably, we observed, exclusively in the human brain, an additional vertical sulcus joining the CGS or PCGS anterior to the rostral part of the genu of the corpus callosum which was labeled the rostro-perpendicular paracingulate sulcus (RP-PCGS) (Fig 7A). In addition, we observed a sulcus located just dorsal to SOS, which joins the lateral surface of the brain but not the CGS or PCGS, called the dorsomedial polar sulcus (DMPS) in 88.75% of hemispheres in human and in 48.75% of hemispheres in chimpanzee. By contrast, only 2.5% of hemispheres of baboon brains display this sulcus and no hemispheres in macaque brains (χ2 = 227.43, df = 3, p-value = 2.2e-16). A second sulcus, the ventromedial polar sulcus (VMPS), located in the ventral medial frontal cortex, joining the lateral surface of the brain and sometimes the ROS-I, was identified more frequently in humans (57.5% of hemispheres) than in chimpanzee (11.25% of hemispheres) and was absent in baboons and rhesus monkeys (χ2 = 126.29, df = 3, p = 2.2e-16) (Fig 7C). Fig 8 summarizes the major changes occurring between rhesus monkeys, baboons, chimpanzees and humans.

**Figure 7.**
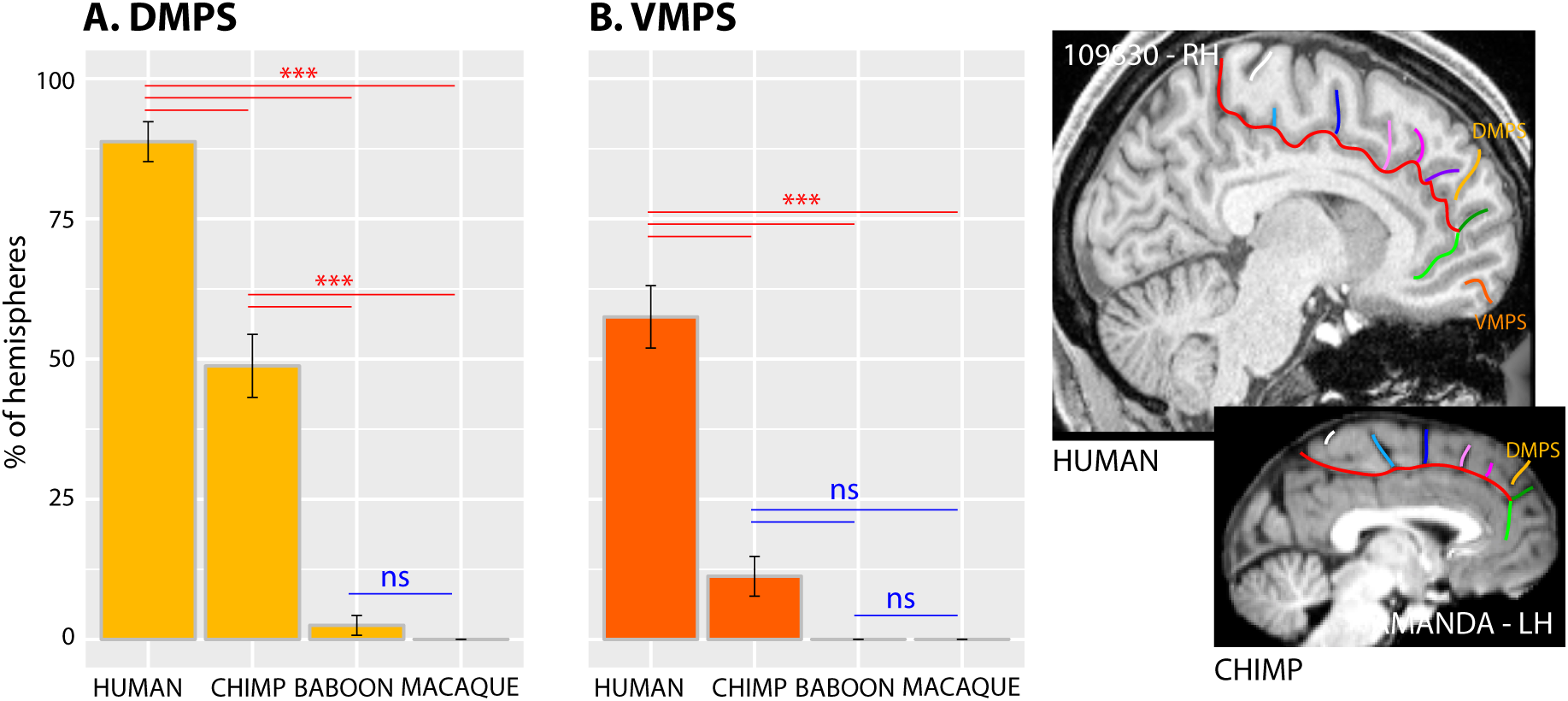
Probability of occurrence of DMPS and VMPS through primates. The location of the DMPS and VMPS sulci are displayed in human and chimpanzee brains in the right panels. The probability of occurrence of both DMPS (**A**) and VMPS (**B**) is higher in human than in chimpanzee but is very low in Cercopithecinae. Statistics: * p<0.05, ** p<0.01, *** p<0.001, ns non-significant.

**Figure 8.**
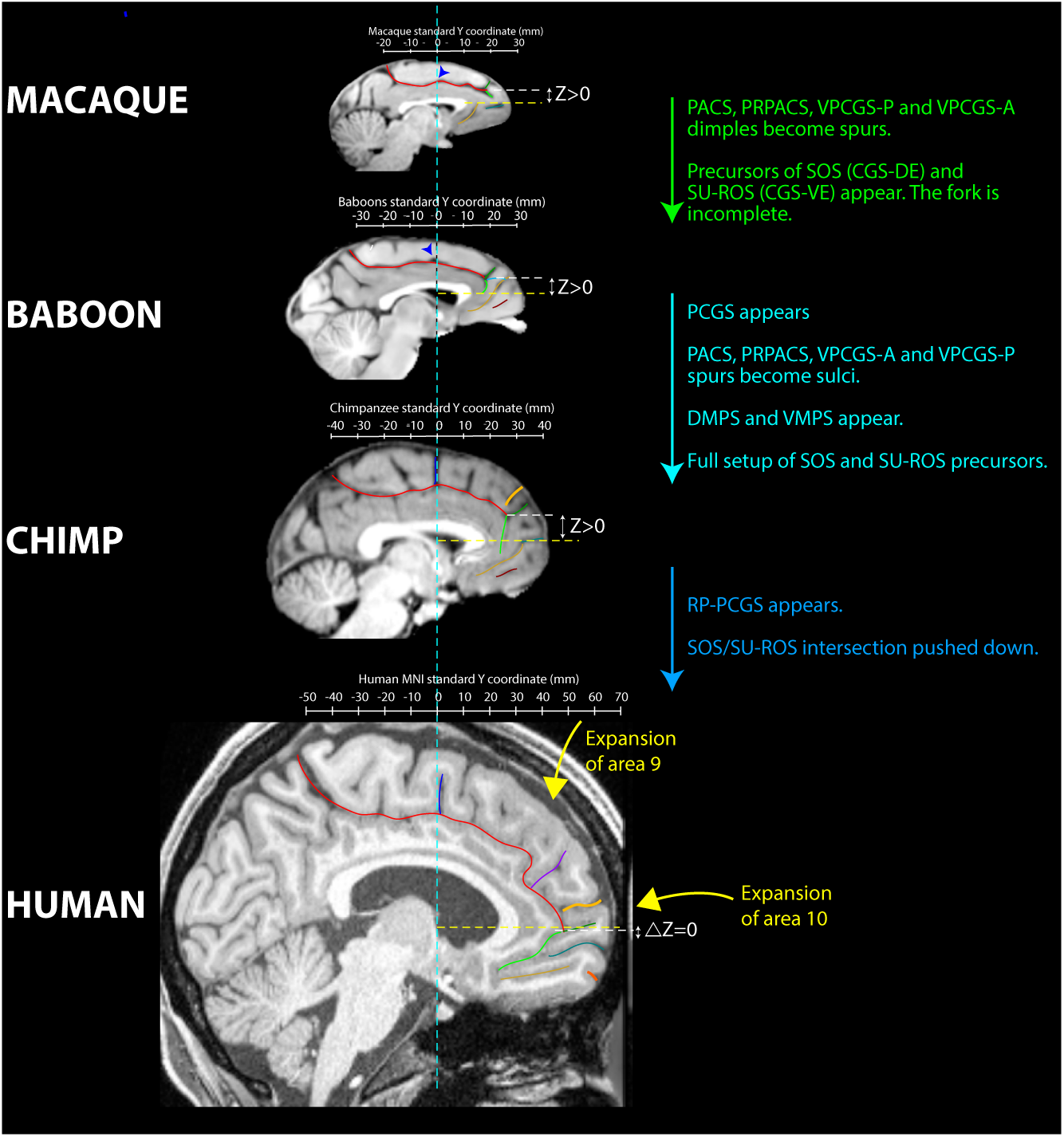
Summary of sulcal changes occurring from macaque to human. Schematic sulcal organization of each specie MFC is represented in their respective stereotaxic space. Main changes from one specie to another are noted on the right. The ΔZ corresponds to the difference between the dorso-ventral Z coordinates where the intersection is observed and the dorso-ventral Z coordinates where the apex of the genu of the corpus callosum is located. It is decreased in human compare to non-human primates, showing that the SU-ROS/SOS and CGS-DE/CGS-VE intersection is pushed down from non-human to human primates.

## DISCUSSION

While gyrification correlates with brain volume (Rogers et al., 2010; Zilles et al., 2013), and therefore with the expansion of primate neocortex, careful analysis of sulcal brain morphology has proven to be a powerful complementary method for a better understanding of the evolution of the medial frontal cortex in primates. Here, we describe for the first time that chimpanzees possess a PCGS, which was previously thought to be a uniquely human feature (Cole et al., 2009, 2010; Vogt, 2016). Two differences must, however, be emphasized: 1) the probability of occurrence of a PCGS is lower in the chimpanzee than in the human brain (present at least in one hemisphere in 70.1% of human versus 33.8% of chimpanzee, Fig 1E), and 2) the presence of a PCGS is largely lateralized in the left hemisphere in the human brain, whereas it is equally present in both hemispheres in the chimpanzee (Fig 1E). Notably, only in the chimpanzee, the probability of occurrence of an ILS is higher in the left than in the right hemisphere (Fig 2A). The ILS being observed only in hemispheres displaying no PCGS in human and chimpanzee (Fig 2B), one can hypothesize that the lateralized presence of an ILS in the left hemisphere in chimpanzee brains may constitute an evolutionary reflection of the beginnings of the sulcal organization as observed in the human brain, i.e. a PCGS lateralized in the left hemisphere and an ILS not lateralized.

Another major difference between the primate species examined was the organization of the medial prefrontal cortex rostral to the genu of the corpus callosum. Differences were associated with 1) a sulcus only observed in humans, the rostro-perpendicular paracingulate sulcus (RP-PCGS), 2) the progressive emergence of the DMPS and VMPS, and 3) the SU-ROS/SOS intersection that is displaced downwards from the level of the apex of the genu of the corpus callosum as one compares more distantly related primates to humans. There is no evidence from cytoarchitectonic (Petrides and Pandya, 1994; Petrides et al., 2012) and neuroimaging (Sallet et al., 2013; Neubert et al., 2015) studies that these changes are associated with new cortical areas, suggesting instead differential expansion of 1) the rostral medial prefrontal cortex (area 9) and medial frontopolar cortex (area 10), two brain regions that have been considered important for high-order socio-cognitive processing, such as mentalizing (Amodio and Frith, 2006; Vaccaro and Fleming, 2018; Wittmann et al., 2018) and 2) the ventromedial frontal cortex (VMPFC) implicated in value-based decisions (Rushworth et al., 2011; Papageorgiou et al., 2017). Our results are in line with a recent study showing that the anterior cingulate/ventromedial prefrontal cortex (ACC/VMPFC) is a region of high structural variability within the human brain and also within the macaque brain (Croxson et al., 2018). By examining the morphological sulcal variability in this region across four primate brains, our results provide an explanation of the nature of this variability.

Furthermore, by comparison with the ACC/VMPFC region, the data demonstrate that, in Old-world monkeys, the organization of the medial frontal cortex (MFC) posterior to the genu of the corpus callosum (where medial motor and premotor areas lie and which corresponds to the Mid-Cingulate Cortex, MCC), has all the basic features of the human sulcal organization and reaches a similar organization across Hominoidea. Specifically, Old world monkeys display precursors –in the form of spurs or dimples-of the four vertical sulci of this region: PACS, PRPACS, VPCGS-P and VPCGS-A. This result is of great importance as it provides additional evidence that the MCC is comparable both anatomically and functionally across macaque and human brains (see Procyk et al. (Procyk et al., 2016) for a detailed anatomo-functional comparison), contrary to what has been proposed by some (Cole et al., 2009, 2010; Vogt, 2016). Importantly, we show for the first time that these sulci refer to specific and highly-reliable anatomical features, and we provide a simple framework to identify these sulci: PACS is located at the level of the rostral limit of the pons, PRPACS at the level of the anterior commissure, VPCGS-P and VPCGS-A at the level of the caudal and rostral limit of the genu of the corpus callosum, respectively. Moreover, these landmarks could be used to guide interpretation of functional neuroimaging studies. Our results also show that the most ventral part of the MFC is highly preserved from macaque to human. ROS-S, which is in the center of this region, is a deep sulcus in all primate species. Importantly, it has been recently shown that the main ventromedial prefrontal node of the default mode network lies in this sulcus in the human brain (Lopez-Persem et al., 2018). Importantly, the macaque default brain network was recently shown to be comparable to that observed in the human brain (Mars et al., 2012; Margulies et al., 2016), providing further support to our present demonstration that the sulcal organization of this region is conserved across the primates examined.

Collectively, the results presented here provide 1) strong evidence for a comparable organization of the MCC across primates and 2) a clear framework to test further hypotheses regarding the relation between the vertical sulci/spurs/dimples, cytoarchitectonic areas, and functions. For example, there is some evidence that PACS may be a sulcus occurring at the border between medial area 4 and area 6 in several primate species: human (Brodmann, 1909; Bucy, 1944; Sarkissov et al., 1955), orangutan (Mauss, 1911; Bucy, 1944), and macaques (Bucy, 1935). Finally, as no new areas have been identified in the human medial frontal cortex in neuroimaging (Sallet et al., 2013; Neubert et al., 2015) and cytoarchitectonic (Petrides and Pandya, 1994; Petrides et al., 2012) studies, one could suggest the emergence of a new sulcus present in the medial cortical region of the human brain might reflect the relative expansion of this region compared to other primates. For instance, we suggest that the RP-PCGS reflects the expansion of medial area 9, a region thought to be important for social metacognitive abilities (Amodio and Frith, 2006; Vaccaro and Fleming, 2018; Wittmann et al., 2018).

Future studies can also test for the relationships between the vertical sulci of the cingulate/paracingulate sulci and cytoarchitectonic areas. Based on cytoarchitectonic studies, Vogt has proposed a hierarchical model of the organization of the cingulate cortex. At the higher level one could distinguish, both in humans and monkeys, a four-region organization (Palomero-Gallagher et al., 2009; Vogt, 2016). This model emphasizes the distinction between the anterior cingulate cortex -ACC- (anterior to the rostral limit of the genu of the corpus callosum), the anterior and the posterior mid-cingulate cortex -aMCC and pMCC-, the posterior cingulate cortex -PCC-, and the retrosplenial cortex. We superimposed this model on our sulcal organization scheme presented here in human and macaque to examine whether the vertical sulci may have significant meanings on the anatomo-functional organization of the cingulate cortex. Based on the consistent location of the vertical sulci we have demonstrated above, one can hypothesize that the VPCGS-A, which is located at the rostral limit of the genu of the corpus callosum and is better conserved than the VPCGS-P across primates, may be a reliable landmark to distinguish the ACC from the aMCC in both human and macaque brains. Vogt’s model identifies the limit between aMCC and pMCC as being the anterior commissure and we demonstrated that the pre-paracentral sulcus (PRPACS) is systematically located at the level of the anterior commissure and is highly conserved across primates. As such, this sulcus might be a limiting sulcus between aMCC and pMCC in both human and macaque brains. Finally, the paracentral sulcus (PACS) which is located at the level of the rostral limit of the pons might be the limit between pMCC and PCC in both human and macaque. Although there is no currently available information about the four-region model in the medial cortex of baboon and chimpanzee brains, our demonstration that the vertical sulci examined are conserved and hold the same location as in human and macaque brains, one can infer that these sulci are limiting the same regions. This hypothesis is displayed in Fig S3 and future studies can assess whether these sulci are limiting functional and cytoarchitectonic areas in these species. Altogether, these data provide strong evidence that the dorsal medial frontal cortex is highly conserved from macaque to human. It is important to note, however, that whether sulci relate to cytoarchitectonic areas remains controversial (Córcoles-Parada et al., 2017; Palomero-Gallagher and Zilles, 2018) and requires considerably more investigations in appropriately sectioned brains. It should be emphasized that proper examination of the relation of sulci to cytoarchitectonic areas requires histological sections perpendicular to the orientation of a given sulcus, as was originally pointed out by Economo and Koskinas (1925) (Economo and Koskinas, 1925).

Besides sulci, we propose that other anatomical landmarks could help guide interpretation of the functional organization of the medial frontal cortex. This is a relevant question as New-world monkeys include species with gyrified brains (e.g. capuchin and squirrel monkeys) and species with lissencephalic brains (e.g. the marmoset) (Van Essen and Dierker, 2007; Chaplin et al., 2013; Donahue et al., 2018). Importantly, it has been shown that the relative position and cytoarchitecture of homologous areas in New versus Old World monkeys are similar (Paxinos et al., 2008, 2012), despite a different brain size (Chaplin et al., 2013). On the basis of the framework provided by the present study, it would be reasonable to hypothesize that homologous cytoarchitectonic and functional regions in the medial frontal cortex in New-world monkeys may be identified using landmarks common to all primates: the PCC/MCC limit should be located at the level of the rostral part of the pons (where PACS is found in old world monkeys, apes, and human). The MCC should expand from this location to the anterior part of the genu of the corpus callosum, i.e. where the VPCGS-A is found and which corresponds to the limit between MCC and ACC. As in humans, we also predict the border between Supplementary Motor Area and Pre-Supplementary Motor Area to be at the level of the anterior commissure in standardized stereotaxic space, i.e. at y=0 in Talairach and Tournoux coordinates (Talairach and Tournoux, 1988) or y=+3 in the latest asymmetrical MNI brain (Collins, 2018).

To conclude, our study provides critical new evidence in the context of a new comprehensive framework regarding prefrontal cortical evolution (summarized in Fig 8). We anticipate that this framework will be used to examine the relationships between sulcal morphology, cytoarchitectonic areal distribution, connectivity, and function, within the studied species but also across the primate order.

## METHODS

Neuroimaging T1 anatomical data of 197 humans, 225 chimpanzees, 88 baboons and 80 macaque brains were analyzed.

### Human subjects

High-resolution anatomical human scans were obtained from the Human Connectome Project database (humanconnectome.org). Only data from subjects displaying no family relationships were analyzed. The participants in the HCP study were recruited from the Missouri Family and Twin Registry that includes individuals born in Missouri (Van Essen et al., 2012). Acquisition parameters of T1 anatomical scans are the following: whole head, 0.7mm^3^ isotropic resolution, TR=2.4s, TE=2.14ms, flip angle = 8° (more details can be found on https://humanconnectome.org/storage/app/media/documentation/s1200/HCP_S1200_Release_Appendix_I.pdf. The full set of inclusion and exclusion criteria is detailed elsewhere (Van Essen et al.,2012). In short, the HCP subjects are healthy individuals who are free from major psychiatric or neurological illnesses. They are drawn from ongoing longitudinal studies (Edens et al., 2010; Sartor et al., 2011; Van Essen et al., 2012), where they received extensive previous assessments including the history of drug use, emotional, and behavioral problems. The experiments were performed in accordance with relevant guidelines and regulations and all experimental protocol was approved by the Institutional Review Board (IRB) (IRB # 201204036; Title: ‘Mapping the Human Connectome: Structure, Function, and Heritability’). All subjects provided written informed consent on forms approved by the Institutional Review Board of Washington University in St Louis. In addition, the present study received approval (n°15-213) from the ethic committee of Inserm (IORG0003254, FWA00005831) and from the Institutional Review Board (IRB00003888) of the French institute of medical research and health.

### Non-human primates

High-resolution anatomical scans of chimpanzee and baboon brains were acquired in the laboratories of Dr. William Hopkins and Dr. Adrien Meguerditchian, respectively. High-resolution anatomical scans of macaque were acquired in the laboratories or Drs E. Procyk/C. Amiez, W. Hopkins, J. Sallet, F. Hadj-Bouziane, and S. Ben Hamed.

### Neuroimaging data analysis

Human and macaque brains were normalized in the human (http://www.bic.mni.mcgill.ca/ServicesAtlases/HomePage) and macaque (Frey et al., 2011) MNI steretotaxic coordinate system, respectively. Chimpanzee brains were normalized in the chimpanzee standard brain developed by Dr. W. Hopkins (Hopkins and Avants, 2013). Baboon brains were normalized in the baboon standard brain developed by Dr. A. Meguerditchian (Love et al., 2016).

Two levels of analysis were performed:

1. **A large qualitative analysis** in the whole set of data (i.e. 197 human, 225 chimpanzee, 88 baboon and 80 macaque brains). This large analysis was performed to identify whether the paracingulate sulcus (PCGS) and the intralimbic sulcus (ILS) were human-specific features. For that goal, we visually inspected each MRI scan to assess the presence or absence of these sulci in both hemispheres of each normalized brain of each species were assessed.
2. **A restricted qualitative/quantitative analysis** of all sulci of the medial frontal cortex in both hemispheres of a subset of 40 brains of each species. This analysis consisted first in assessing the characteristics of the vertical sulci in the superior medial frontal cortex emerging from the cingulate (CGS) or the PCGS in all brains across species. For that goal, the sulci were assigned in one of 3 categories based on our qualitative observations (Fig S1): <u>sulcus</u> (a real sulcus -long and deep- can be observed), <u>spur</u> (a precursor sulcus, i.e. not long enough to be considered a sulcus), and <u>dimple</u> (locations where a hump indicates a dimple of the CGS).

Second, this analysis consisted in identifying all sulci in the medial frontal cortex, all anatomical landmarks fixed across species (rostral limit of the pons, anterior commissure, caudal and rostral limit of the genu of the corpus callosum, apex of the genu of the corpus callosum) and in identifying any relationships that may exist between the location of a given sulcus and a given anatomical landmark in all species. The relationships between the location of a given sulcus and a given anatomical landmark in all species was performed as follow:

a. To assess whether PACS, PRPACS, VPCGS-P, and VPCGS-A were located at the level of, respectively, the most rostral limit of the pons (landmark 1), the anterior commissure (landmark 2), the caudal (landmark 3) and the rostral (landmark 4) limit of the genu of the corpus callosum, we calculated the difference between the Y value of the intersection between the CGS or the PCGS (if present) and the PACS, PRPACS, VPCGS-P, and VPCGS-A and the Y value of landmarks 1, 2, 3, and 4, respectively. This difference was calculated in all species on the normalized T1 data of the four species. This difference was then normalized to take into account the different antero-posterior extent of the brains of the four species (i.e. antero-posterior extent of the human, chimpanzee, baboon, and macaque brains are respectively 175, 110, 85 and 60 mm).
b. To assess the relative location of the intersection of the fork formed by SU-ROS/CGS-VE and SOS/CGS-DE with the CGS or PCGS across primates, we calculated the % of displacement of this intersection from the apex of the genu of the corpus callosum. In that goal, we measured the difference between the Z value of this intersection and the Z value of the apex of the genu of the corpus callosum. To compare primate brains, this difference was then normalized on the basis of the dorso-ventral extent of the brains normalized in their respective standard space (see Methods) at the level of the genu of the corpus callous across primates (i.e. 100, 50, 30, and 24mm, respectively in human, chimpanzee, baboon, and macaque brains).

### Statistical analysis

Concerning binomial data, we built binomial logistic regression GLMs. ANOVA Chi-square tests and post-hoc Tukey tests were then applied. Note that, in the cases where only one value was observed in a specific variable (e.g. the PCGS is absent in baboon and macaque and the “presence” value is therefore 0 for all subjects), the binomial logistic regression GLM was fitted using an adjusted-score approach to bias reduction (using the brglm package (https://cran.r-project.org/web/packages/brglm/brglm.pdf).

Concerning gaussian data (e.g. displacement of the fork formed by SU-ROS/CGS-VE and SOS/CGS-DE), regular GLM analyses, ANOVA F tests, and Post-hoc Tukey tests were applied to assess whether the location of the various sulci. All statistics were performed with R software, R Development Core Team (R, 2008) under R-Studio (R, 2016).

## Acknowledgments

This work was supported by the Human Frontier Science Program (RGP0044/2014), the Canadian Institutes of Health Research (CIHR) Foundation grant FDN-143212, the Medical Research Foundation (FRM), the Neurodis Foundation, the French National Research Agency, and the labex CORTEX ANR-11-LABX-0042 of Université de Lyon. E.P. and C.A. are employed by the Centre National de la Recherche Scientifique. J.S. was supported by a Sir Henry Dale Wellcome Trust Fellowship (105651/Z/14/Z). The Wellcome Centre for Integrative Neuroimaging is supported by core funding from the Wellcome Trust (203139/Z/16/Z). We thank K. Knoblauch for helpful comments on statistical analysis. We thank Delphine Autran-Clavagnier for data acquisition in macaques.

Author Contribution
C.A. built the project, coordinated the consortium to gather the dataset, analyzed and interpreted data, and wrote the article. J.S. and M.P. interpreted data and wrote the article. W.D.H. provided chimpanzee and macaque T1 data, A.M. provided baboon T1 data, F.H-B., S.B., C.R.E., and E.P. provided macaque T1 data.

