## Supplementary material for "Sulcal organization in the medial frontal cortex reveals insights into primate brain evolution": Fig S3

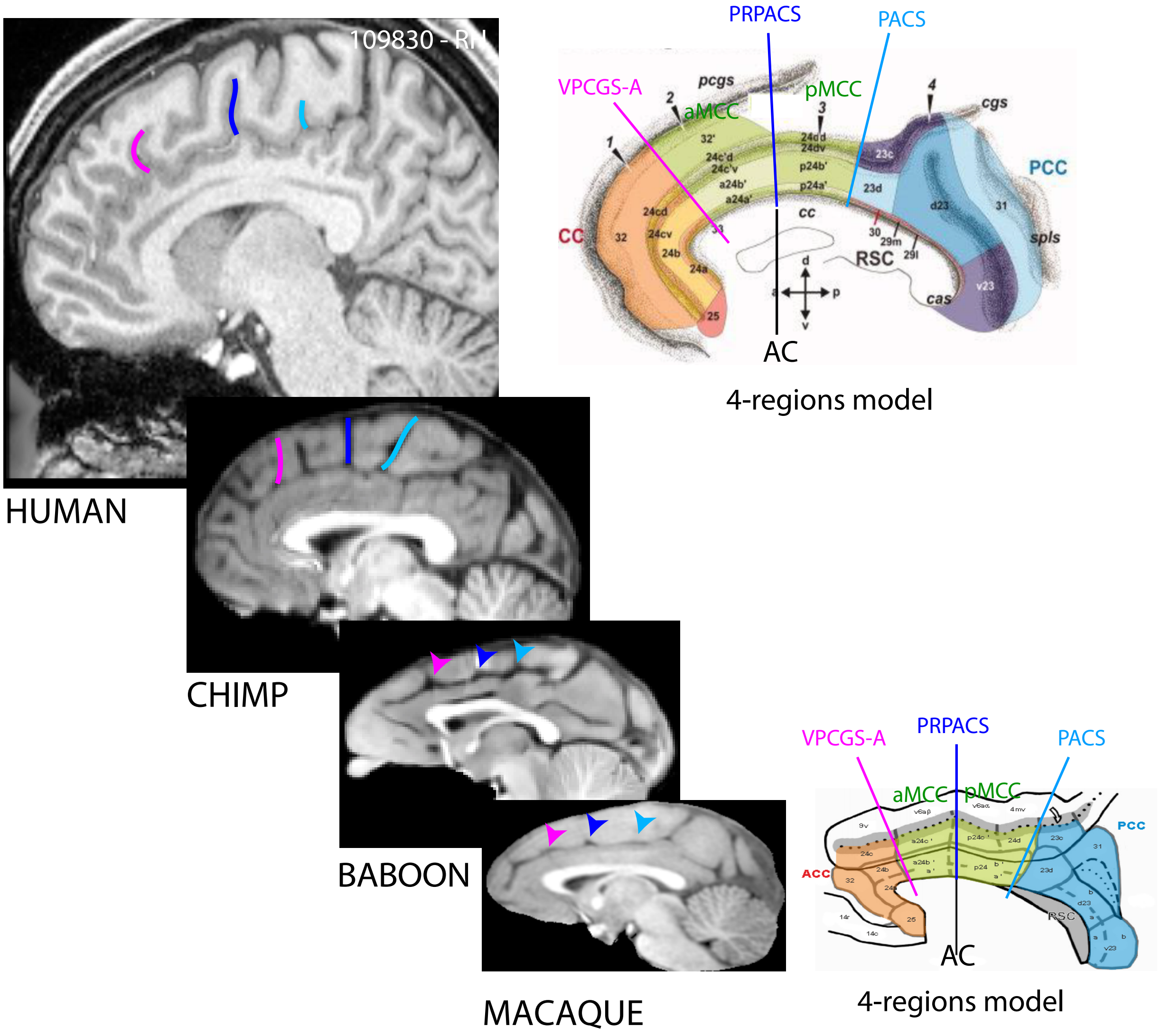


Figure S3. Possible relationships between the vertical sulci in human and macaque and cytoarchitectonic areas in the MCC from the 4-regions model developed in both species 42,76. One hypothesis would be that the A-VPCGS is the limiting sulcus between the ACC and the aMCC in both human and macaque. Vogt's model identifies the limit between aMCC and pMCC as being the anterior commissure. PRPACS is systematically located at the level of the anterior commissure and is conserved across primates. As such, this sulcus might be a limiting sulcus between aMCC and pMCC in both human and macaque. Finally, PACS being located at the level of the rostral limit of the pons, it might correspond to the limit between pMCC and PCC in both human and macaque. These three vertical sulci being highly preserved in baboon and in chimpanzee and displaying the same location than in human and macaque, one can infer that these sulci are limiting the same areas than in human and macaque.
