## Supplementary material for "Sulcal organization in the medial frontal cortex reveals insights into primate brain evolution": Fig S2

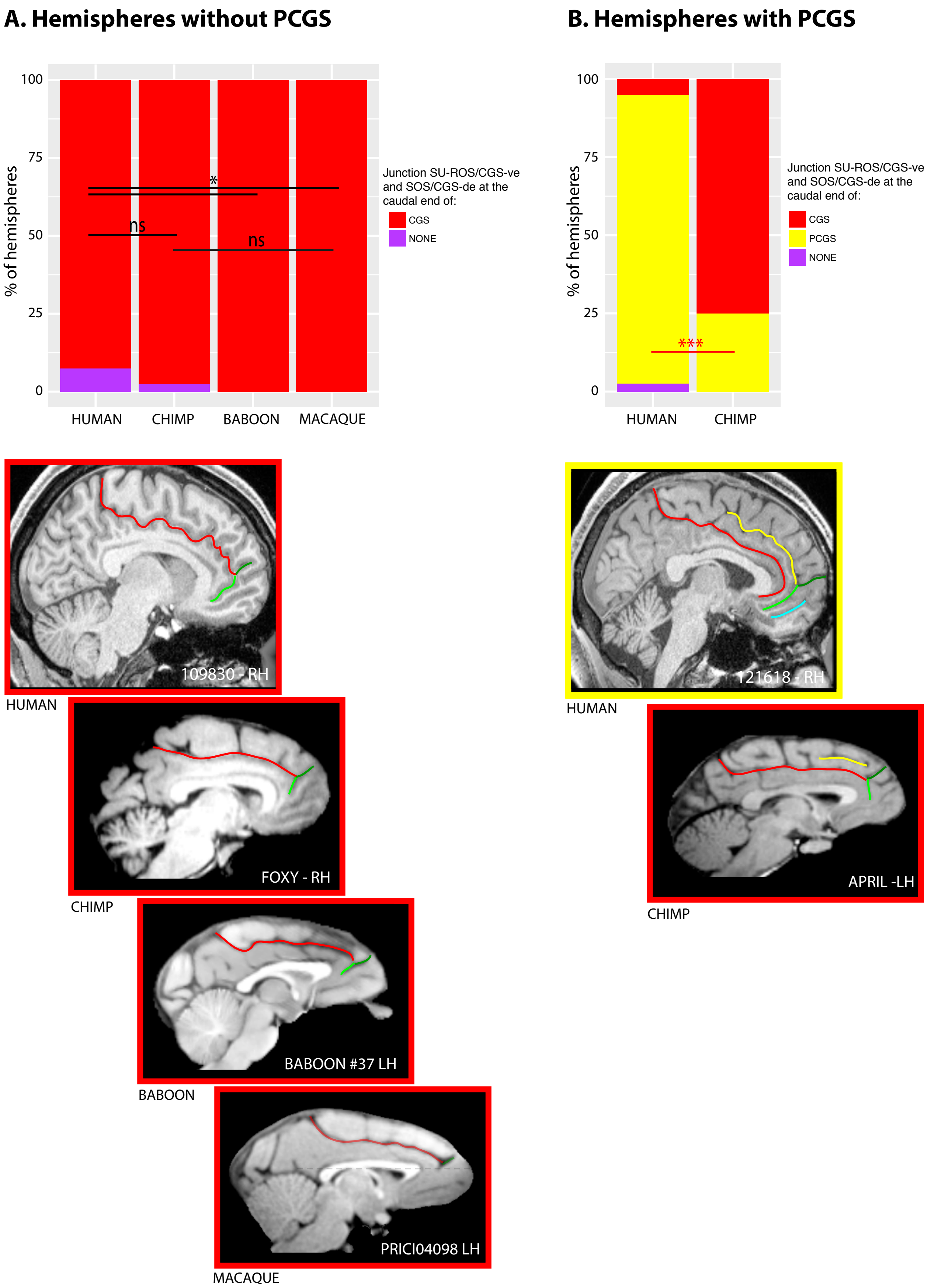


Figure S2. Location of the SU-ROS/SOS and CGS-DE/CGS-VE intersection in hemispheres without (A) and with a PCGS (B) in primates. In human, when the PCGS is absent, the intersection is located at the rostral end of the CGS but when the PCGS is present, it is located at the rostral end of the PCGS in the large majority of hemispheres. In chimpanzee, when the PCGS is absent, the intersection is located at the rostral end of the CGS. By contrast, when the PCGS is present, it is still located at the rostral end of the CGS -and not of the PCGS as in human- in the large majority of hemispheres. In baboon and macaque, the intersection is located at the rostral end of the CGS.
