## Supplementary material for "Sulcal organization in the medial frontal cortex reveals insights into primate brain evolution": Fig S1

### VERTICAL SULCI DESCRIPTION

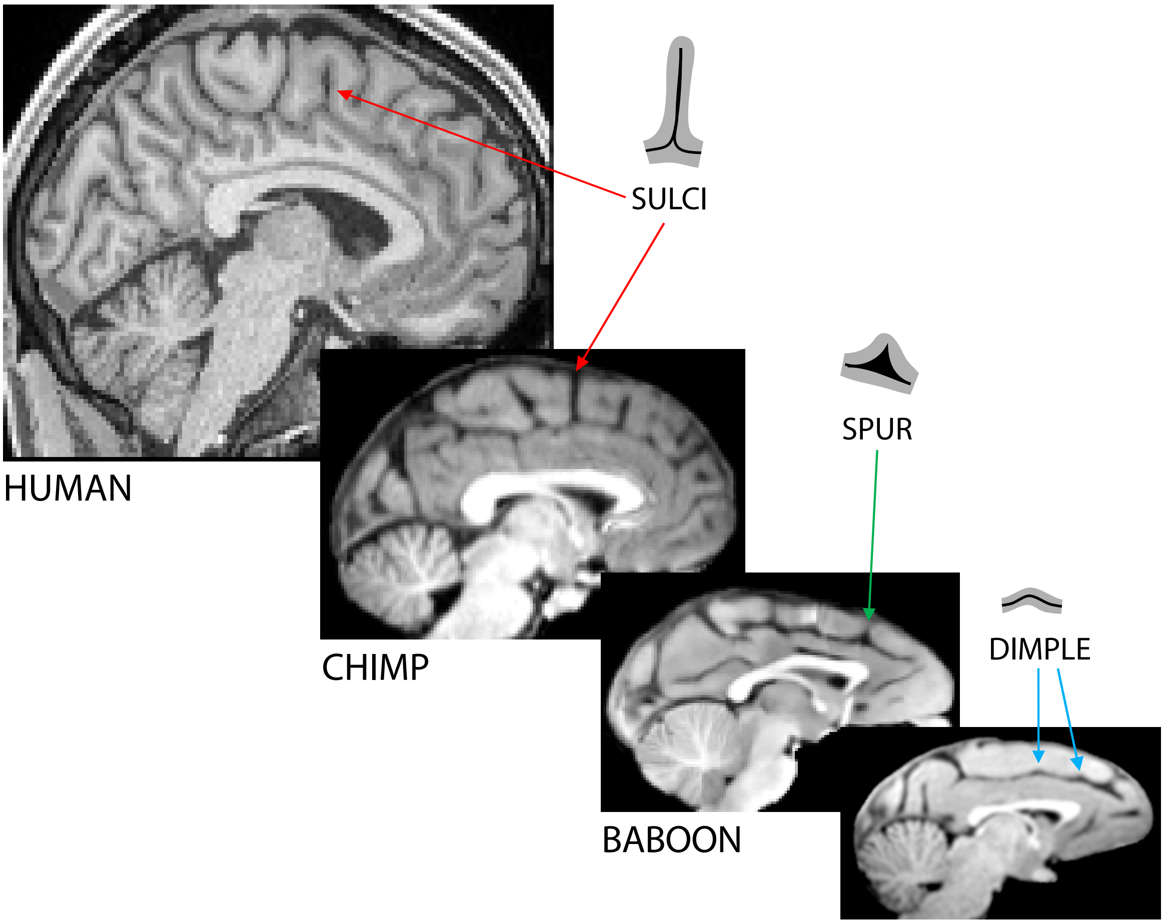


Figure S1. Description of the vertical sulci in the MCC. Vertical sulci are deep sulci in human and chimpanzee, spurs or dimples in baboons.
